# Diet-dependent microbiota and diet-independent immunometabolic responses to probiotic supplementation in broiler chickens

**DOI:** 10.64898/2026.06.08.730860

**Authors:** Lauren E. Anderson, Anne L. Ballou, Natalie B. Roberts, Rizwana A. Ali, Matthew D. Koci

## Abstract

Probiotics are widely used in food animal production to support gut health and immune function, but the indicators of probiotic efficacy and the conditions under which they translate to host benefit remain unclear. Microbiota composition is the most accessible data supporting probiotic effects, yet whether compositional change reliably predicts host outcomes is not well understood. We investigated this question in broiler chickens fed two nutritionally similar basal diets, with or without a commercial probiotic. Microbiota composition was profiled across 6 gastrointestinal regions using 16S rRNA sequencing. To assess systemic functional effects, an *in vitro* assay building on prior observations of elevated circulating immune cell ATP in probiotic-fed animals was developed. In this assay, serum from each treatment group was applied to a chicken T-lymphocyte cell line before ATP quantitation. Basal diet was the primary driver of microbial community structure, with probiotic-induced compositional shifts observed predominantly in one diet context but minimally in the other. Despite this difference, serum from probiotic-supplemented animals increased T-lymphocyte ATP production across both diets, supporting prior findings and revealing a systemic immunometabolic response independent of broad microbiota restructuring. Functional predictions revealed enrichment of pathways related to mevalonate and carbohydrate metabolism in probiotic-supplemented birds within the more responsive diet context, driven largely by Lactobacillaceae family taxa. These findings demonstrate that basal diet modulates the detectability and nature of probiotic effects on the microbiota, but not the physiological host response. This disconnect has implications for how probiotic efficacy is evaluated and for microbiome targeted interventions across species.

**Importance:** Probiotics are used widely in food animal production to support gut health and immune function, yet predicting which probiotic preparations will produce meaningful effects remains a challenge. Microbiota composition, profiled by 16S rRNA sequencing, is the most accessible measure of probiotic activity, but it captures only one aspect of the host-microbe dynamic. These data demonstrate that probiotic-induced compositional changes vary substantially between basal diets, while the host immunometabolic response is consistent across diets, demonstrating that compositional readouts alone cannot reliably predict host outcomes. The findings have practical implications for how probiotic efficacy is evaluated and inform the broader effort to design microbiome targeted interventions across both veterinary and human contexts.

## Introduction

Probiotics are widely used in food animal production to support gut health, immune development, and overall performance (1, 2). However, the mechanisms of action and the indicators that reliably predict probiotic efficacy remain unclear. This likely reflects context dependent interactions between probiotic strains and resident gut microbiota shaped by host species, production environments, diets, housing conditions, microbial exposure, etc. (3, 4). These factors influence microbial community assembly and function, yet few studies evaluate how they affect probiotic activity along the entire gastrointestinal (GI) tract or how those changes connect to host outcomes. This gap limits the design and evaluation of probiotic interventions in both veterinary and human contexts.

Even when diets are formulated to meet comparable nutritional specifications, the ingredients used to achieve those targets can vary. Regional sourcing, storage conditions, and processing practices contribute to shifts in feed composition and microbial content (5, 6). Diet form also affects mechanical digestion and chemical conditions in the upper GI tract, which in turn shape microbial niches downstream. Together, these aspects define the microbial environment into which probiotics are introduced and influence the extent to which microbial communities shift in ways observable by sequence-based methods.

Our previous studies have demonstrated that specific probiotic supplementation can alter immune function and energetics, both in the local intestine level and systemically. Over multiple studies, we observed decreased expression of pro-inflammatory cytokines in the small intestine, increased energy consumption by the thymus and circulating peripheral blood mononuclear cells (PBMCs), as well as a more rapid immune response to challenge in probiotic-fed broilers (7, 8). These findings suggest that probiotic exposure shifts immunometabolism but did not address whether these effects related to specific changes in the gut microbial community.

To address this gap, we evaluated how two nutritionally similar broiler starter diets, with different ingredient sourcing and physical structure, shaped the microbiota and host response to probiotic supplementation. Microbial communities were profiled across 6 gut regions using 16S rRNA sequencing. To assess systemic immunometabolic effects, we developed an *in vitro* assay in which pooled serum from each treatment group was applied to a chicken T-lymphocyte cell line, allowing evaluation of whether microbiome-derived serum compounds could influence immune cell ATP production *in vitro*.

By combining spatial microbiome profiling with a functional assessment of immune cell metabolism, this study demonstrates how diet shapes the microbial response to probiotic supplementation while the systemic immunometabolic response is preserved across diets. This indicates that differences in the probiotic effect within diet do not necessarily translate to differences in physiological outcomes.

## Methods

### Animal Trial and Diet Formulation

Twenty-four one-day-old, mixed-sex, broiler-type chicks were randomly assigned to one of 4 treatment groups (n = 6 per group) in a 2 × 2 factorial design: two diet types with probiotic supplementation (PB) or without (CON). The mash diet was produced at North Carolina State University Feed Mill Education Unit (MD), while the crumbled diet was a commercially available diet (CD) purchased from LabDiet® (St. Louis, MO). Both diets were nutritionally complete and compositionally similar, formulated to meet or exceed NRC recommendations (Supplemental Table 1), and devoid of any antibiotics. The PB treatment contained 0.3% (w/w) PrimaLac® (Star Labs Inc., Clarksdale, MO), a blend of *Lactobacillus acidophilus*, *Lacticaseibacillus casei*, *Enterococcus faecium*, and *Bifidobacterium bifidum* with a rice hull meal carrier (complete taxa profile in Supplemental Figure 1). The probiotic was added to the feed after processing to prevent thermal degradation. This resulted in approximately 3 × 10⁵ CFU/gram of feed. Birds in all treatment groups were fed *ad libitum* for 4 weeks. All animal procedures were approved by the North Carolina State University Institutional Animal Care and Use Committee (OLAW #A3331 - 01).

### Sample Collection

On day 28, animals were euthanized, and serum was collected, as well as digesta samples from 6 regions of the GI tract: crop, gizzard, duodenum, jejunum, ileum, and cecum. The contents of the crop was gently mixed by hand before sampling. The entire gizzard contents were collected due to low volume. For each section of the small intestine, digesta samples were collected from the midpoint. Cecal content was collected from one cecum per bird. All samples were snap-frozen in liquid nitrogen and stored at -80°C.

### T-lymphocyte ATP Production Assay

To achieve sufficient volume, serum from each treatment group was pooled and used to supplement culture media for Cu205 T-lymphocyte cell line (9). Cells were counted using a hemocytometer, seeded at 5,000 cells/well in RPMI media supplemented with 1% heat-inactivated fetal bovine serum, 1% L-glutamine, and 10% heat-inactivated chicken serum (from CD+CON, CD+PB, MD+CON, or MD+PB groups). Cells were incubated for 48 hours at 41°C and 6% CO₂, then reseeded at 5,000 cells/well in 96-well plates and incubated for an additional 48 hours. ATP was then quantified using an ATP-dependent luciferase assay (CellTiterGlo; Promega, Madison, WI) and compared to an ATP standard curve. Bioluminescence was measured using a Thermo Fluoroskan Ascent FL (Thermo Scientific, Waltham, MA). ATP was normalized to cell number to account for any differences in proliferation using replicate wells with an intercalculating DNA dye (CyQuant NF; Thermo Scientific, Waltham, MA) and a Thermo Fluoroskan Ascent FL at 485/520nm ex/em. Model residuals were approximately normally distributed according to a Shapiro-Wilk test (p = 0.07), but variances differed significantly between groups via Levene’s test (p = 0.0003). Therefore, a heteroscedasticity-robust (HC3), 2-way linear model was used to test the effect of Diet, Treatment, and their interaction with Tukey post-hoc pairwise comparisons.

### DNA Extraction and 16S rRNA Sequencing

DNA was extracted from digesta using the MO BIO PowerSoil kit (MO BIO, Carlsbad, CA) with minor protocol modifications to enhance microbial lysis. Samples were heat-treated at 65°C for 10 minutes, then homogenized in garnet bead tubes at 5,100 RPM for 45 seconds using a Precellys 24 homogenizer. DNA concentrations were quantified using a NanoDrop 2000 spectrophotometer and normalized prior to sequencing. DNA was also extracted from the PB supplement, as well as each of the 4 diets, using the same protocol to assess microbial presence in the diet and the impact of PB inclusion.

Amplicons targeting the V4 region of the 16S rRNA gene were generated using the Earth Microbiome Project primer set (515F/806R) and sequenced on an Illumina MiSeq platform (151 × 151 bp paired-end) at the Argonne National Laboratory.

### Sequence Data Analysis

The unpaired, raw sequencing reads were paired and filtered using EA-Utils33 (10). Paired reads were processed using the QIIME2 suite of tools (11), barcode matching and Deblur (12) quality filtering (median quality score 33) was conducted prior to assigning amplicon sequence variants (ASVs) and classification using the SILVA database (v 138.1) (13, 14). One CD+CON duodenum sample (−3.05 standard deviations from the mean) and one CD+PB ileum sample (+6.57 standard deviations from the mean) were excluded from analysis due to excessive deviation from the mean sequencing depth (Mean: 11,471; Standard Deviation: 3,842). Three samples (MD+PB ileum, CD+PB duodenum, and CD+CON ileum) were removed from the study due to potential cecal contamination. Mitochondria and chloroplast features were filtered from the remaining samples before analysis. Sequencing of feed samples indicated that the majority of sequencing reads (98% in CD and 99% in MD) were classified as chloroplasts. Sequences were submitted to the Sequence Read Archive at the BioProject accession number PRJNA1450620.

Data were transformed using Center Log Ratio (CLR) in the Phyloseq (15) package in R, and Principal components analysis (PCA; with taxa loadings) were created using the MicroViz package (16). In cases where ASVs were not identified within all samples, read counts with a zero value were managed using a pseudocount equaling one half of the minimum non-zero value for that ASV in order to apply CLR (17).

Phylogenetic Investigation of Communities by Reconstruction of Unobserved States (PICRUSt2) (18) was used to map ASVs to metagenomic reference sequences. Predicted pathway abundances within each sample were inferred using the MetaCyc (19) database. Differential abundances of taxa and MetaCyc inferred pathways were determined using Microbiome Multivariable Associations with Linear Models (MaAsLin2) (20). To assess basal diet effect, regardless of PB supplementation, MD and CD groups were compared to each other with PB as a covariate. A Benjamini-Hochberg false discovery rate (FDR) adjustment was applied to correct for multiple testing, and an adjusted value ≤ 0.1 was considered significantly different. Other taxa and PICRUSt2 differential abundance and ASV contributions visualizations were generated using Tableau (21).

## Results and Discussion

### PB-mediated changes in feed microbial composition differed between basal diets

To identify microbial compositional differences in the 4 diets, 16S rRNA analysis of the PB supplement and each diet+treatment combination was performed in duplicate for a quality control (QC) screen. Because the dataset consisted of duplicates, abundance differences could not be statistically evaluated, and we have reported qualitative observations only. After chloroplast removal, the PB supplement contained 74 different genera. The genus *Pantoea*, often found on grain crops (22), was the most abundant taxa in the PB supplement and appeared at a greater relative abundance in both MD+PB and CD+PB feed compared to CON (Figure 1A). A similar pattern was observed in an unclassified genus within the Lactobacillaceae family. It was one of the most abundant genera in the PB supplement and was exclusively detected in PB-supplemented feed from both MD and CD diet types (Figure 1A, Figure 1B). However, despite its exclusivity to PB feed, this unclassified Lacotobacillaceae ASV was not exclusive to PB-treated animals’ GI tracts (Figure 1B), indicating that its acquisition does not depend on feed.

**Figure 1.**
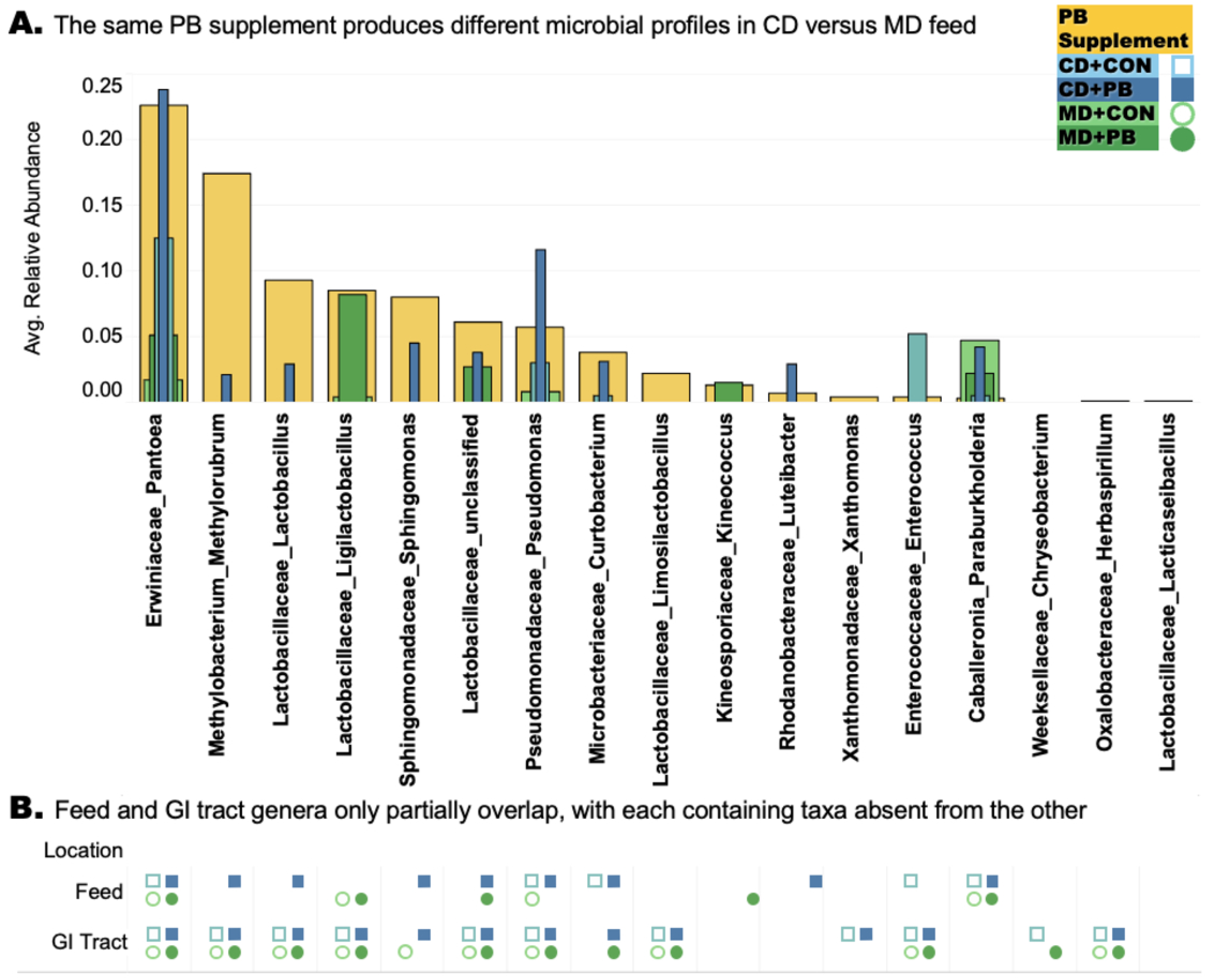
Diet context and GI tract environment influence which PB-associated genera appear in feed and persist in the gut. Samples of each feed type (n = 2/type) and the probiotic supplement (n = 2) were subjected to 16S rRNA sequencing. **(A)** Average relative abundance of bacterial genera in the probiotic supplement (yellow), basal MD (light green) and CD (light blue), and PB treated MD (dark green) and CD (dark blue) are depicted as layered bars. **(B)** The presence or absence of these genera in the feed or gastrointestinal tract (crop, gizzard, duodenum, jejunum, ileum, cecum) of animals fed MD (green, circles) or CD (blue, squares) diet, each with (+PB; darker color, filled shapes) or without (CON; lighter color, open shapes) a commercial probiotic. Colors represent diet and treatment as in panel A. Taxa shown here were selected to support interpretive clarity. Complete data are available in Supplemental Figure 1.

There were also PB effects that differed between CD and MD diets, despite both receiving the same PB treatment. For example, *Pseudomonas* appeared to be more abundant in CD+PB than CD+CON (Figure 1A), but it was not detected in MD+PB feed at all (though present in MD+CON; Figure 1B). *Ligilactobacillus* appeared to be more abundant in MD+PB feed compared to MD+CON (Figure 1A) but was not present in either CD diet (Figure 1B). *Curtobacterium* appeared to be more abundant in CD+PB feed than CD+CON (Figure 1A), but it did not appear in MD diets at all (Figure 1B). Four additional genera present in the PB supplement appeared exclusively in CD+PB feed (e.g., *Methylorubrum*, *Lactobacillus*, *Sphingomonas*, and *Luteibacter*), while only 1 was exclusively found in MD+PB (*Kineococcus*; Figure 1A & B). The presence-absence and qualitative abundance patterns suggest that PB-associated microbes may have been more detectable in CD than in MD, potentially due to diet-specific ingredient composition or basal microbial load, as many PB-associated genera existed in CON diets prior to the addition of PB.

Despite these differences in feed microbial composition, some PB or feed-associated genera were not subsequently detected in the GI tract, including *Kineococcus* and *Luteibacter* (Figure 1B). This lack of GI detection could reflect limited colonization, presence below detection thresholds, or transient functional effects that do not require persistent colonization (23, 24). Additionally, some members of the GI microbiota may be shaped by host selection pressures rather than direct microbial transfer from feed, as they were detected in the GI but not in feed (Figure 2B).

**Figure 2.**
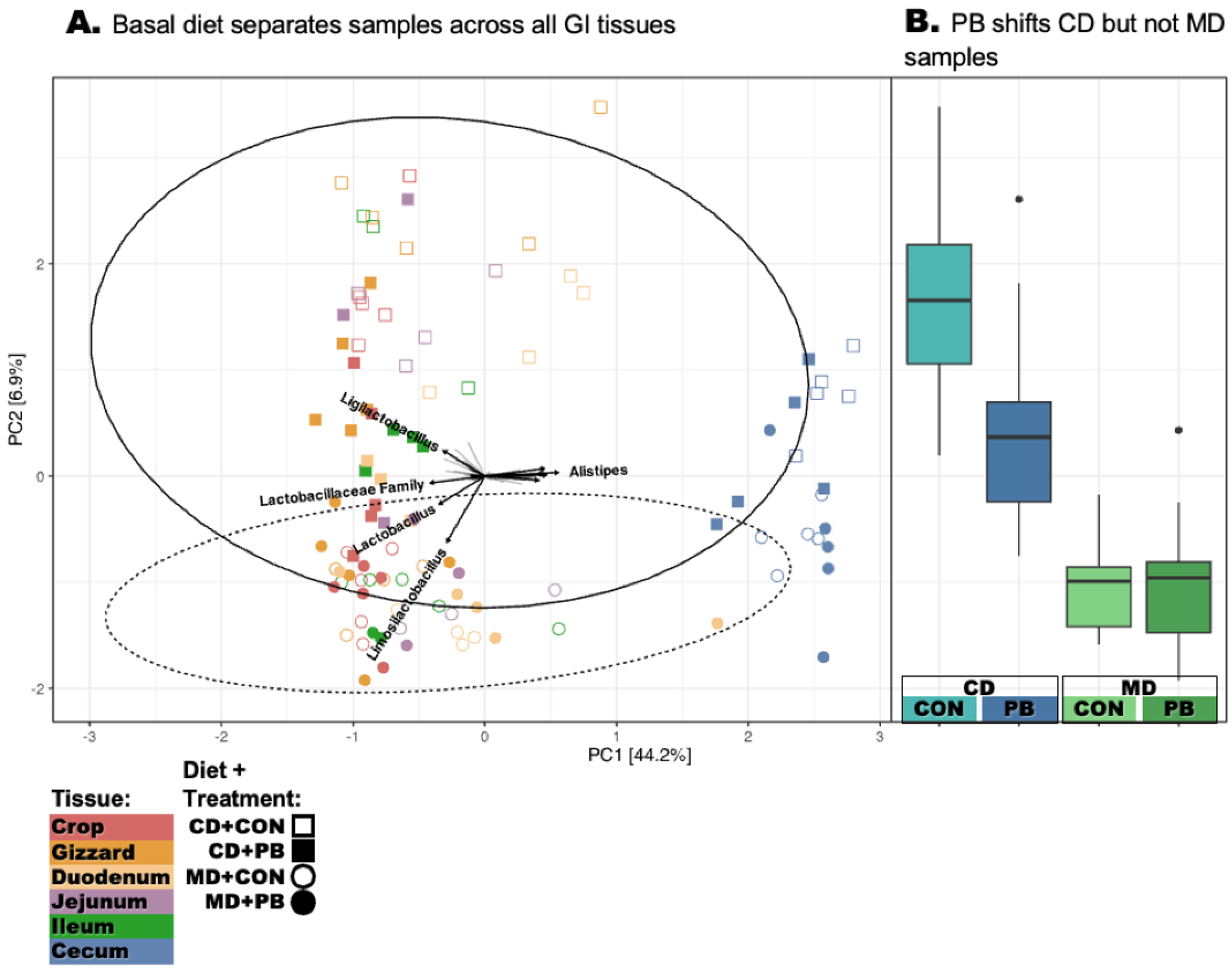
PB alters microbial community structure in CD-fed but not MD-fed birds. **(A)** Principal components analysis (PCA) of center log transformed (CLR) abundance data for all tissue types (Crop=Red, Gizzard=Orange, Duodenum=Yellow, Jejunum=Purple, Ileum=Green, Cecum=Blue), Diets (MD=Squares, CD=Circles), and Treatments (CON=Open Shapes, PB=Filled Shapes) was performed, with genus-level loadings overlaid to highlight key contributors to variation. Ellipses illustrate separation by basal diet, with CD and MD forming distinct clusters. **(B)** Boxplots illustrate PB-induced separation within diet on PC2.

A more robust, replicated study beyond this initial QC screen will be required to determine interactions between feed microbiomes and probiotics. Though these exploratory results suggest that CD feed may have offered a more permissive community for PB-associated taxa to persist or remain detectable, and feed microbial composition alone did not determine downstream colonization in terms of presence-absence. However, the broader community context within the GI tract may provide a more informative view of how the basal diet and PB supplement influence microbial ecology.

### GI microbial communities were primarily structured by basal diet, with greater PB compositional responsiveness in CD

Microbial community structure varied by GI region, consistent with physiological and environmental gradients along the GI tract (6, 25) (Supplemental Figure 2). PCA clustering of CLR transformed abundance data revealed that tissue type was the primary driver of microbiota variation, with basal diet as a secondary influence (Figure 2A). Within this space, MD samples were enriched for *Limosilactobacillus* while *Ligilactobacillus* was strongly associated with CD samples (Figure 2A), despite it only appearing in MD feed (Figure 1B), reinforcing that diet shapes microbiota through mechanisms beyond simple feed-to-gut colonization. In contrast to diet effects, PB effects were limited and interacted with dietary context (Figure 2 A-B). Within the CD diet, samples separated by PB treatment, however, PB supplementation did not alter microbial community clustering in MD (Figure 2B).

It is well understood that diets strongly shape the GI microbiome (26, 27), so differences between MD and CD communities were anticipated. However, it was unexpected that PB-induced shifts in community structure were only observed in CD. This raised the possibility that PB-induced physiological outcomes may also be diet-dependent.

### Pooled serum from PB-supplemented animals reproducibly increased lymphocyte ATP production across both basal diets

Prior findings indicated that PBMCs isolated from PB-fed animals had enhanced ATP production (8). In order to interrogate systemic impacts of PB supplementation on immune energetics, an *in vitro* assay was developed in which the Cu205 T-lymphocyte cell line was treated with serum from MD+CON, MD+PB, CD+CON, or CD+PB chickens and then assayed for differences in ATP concentrations. Robust 2-way analysis indicated significant main effects of diet (p = 0.007) and treatment (p=0.0001), but no significant diet by treatment interaction (p = 0.22). Despite the limited microbial response to PB supplementation in MD-fed animals, serum from PB-treated animals increased Cu205 ATP production compared to CON in both MD and CD diet contexts (Figure 3). ATP levels were lower in MD samples compared to CD, indicating that diet alone establishes baseline energetic differences before PB exposure. Because serum was pooled, this assay does not capture inter-bird variability, but replicate assays using the same pooled serum demonstrated reproducibility and directionality of PB-associated effects.

**Figure 3.**
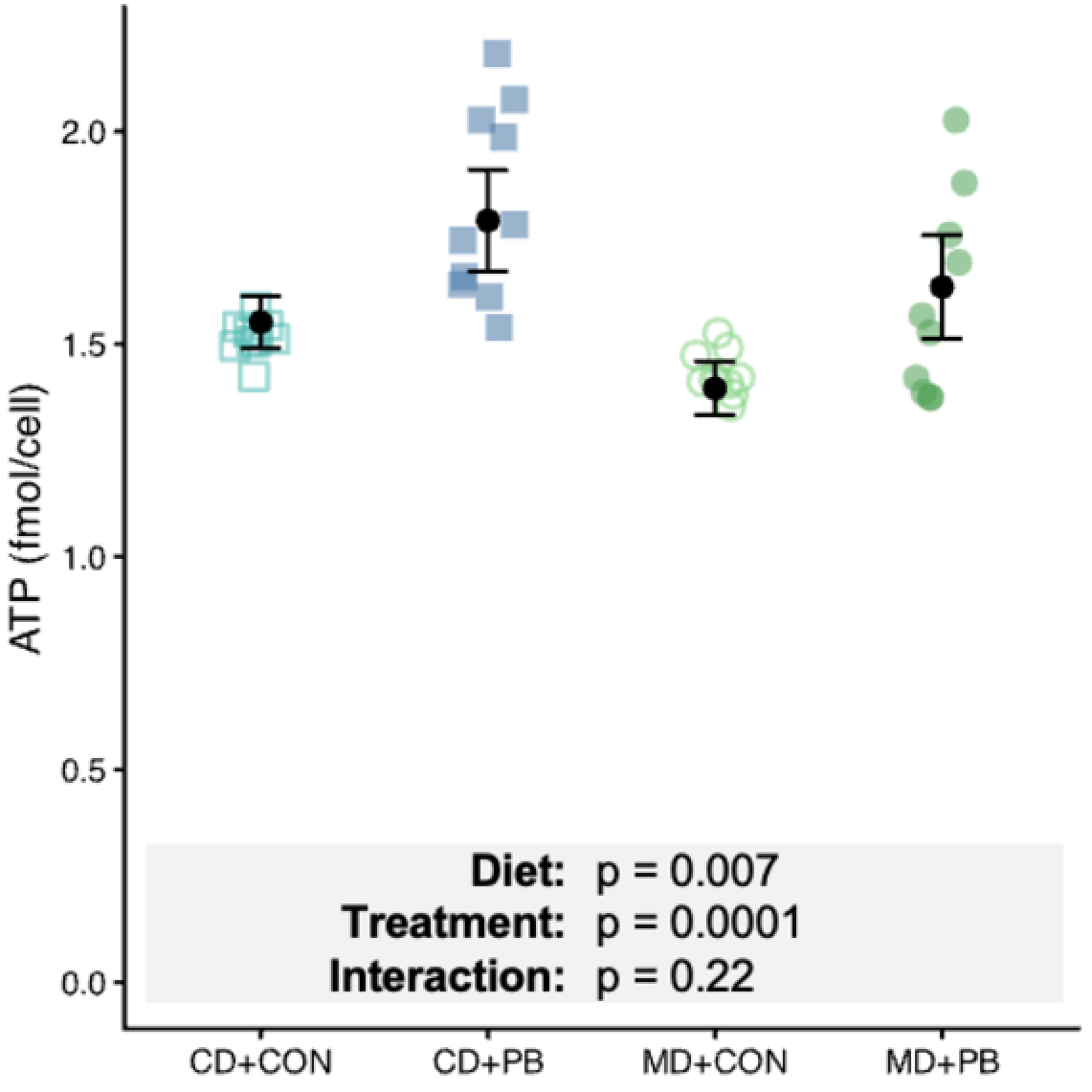
Pooled serum from probiotic-supplemented birds reproducibly increased ATP production in cultured T-lymphocytes across both basal diet types. A chicken T-lymphocyte-like cell line, Cu205, was cultured in media supplemented with 10% pooled serum collected from CD+CON, CD+PB, MD+CON, or MD+PB fed chickens. The original pooled serum from each treatment group was evaluated across repeated assay runs conducted over multiple days. ATP/cell was quantified using a luciferase assay and normalized to cell number determined by nuclear fluorescence. ATP concentration was calculated from a 2-fold standard curve (0.250 nmol/well), and cell counts were verified by serial dilution of Cu205 cells counted with a hemocytometer. Points represent individual measurements from pooled serum samples. Black points and error bars indicate least-squares means with 95% confidence intervals from a heteroscedasticity-robust linear model (HC3).

The observation that PB serum increased lymphocyte ATP production in both diet contexts, despite the lack of detectable PB-associated clustering shifts within MD, indicates that physiologically meaningful changes may occur without overt restructuring of the entire microbiota and prompts examination at a finer taxonomic resolution.

### PB-induced microbial shifts were taxonomically distinct in CD-fed animals

Differential abundance analysis revealed both genus level (Figure 4A) and ASV level (Figure 4B) differences attributable to basal diet alone (Figure A1, Figure B1), as well as PB supplementation under the CD diet (Figure A2, Figure B2). PB-associated changes were limited in MD-fed animals, as might be expected given the lack of PCA separation (Figure 2). There were only 3 ASVs differentially abundant between MD+CON and MD+PB: 2 unclassified ASVs that decreased with PB treatment, and an ASV within the *Faecalibacterium* genus increased (Supplemental Table 2), but there were no differences at the genus level.

**Figure 4.**
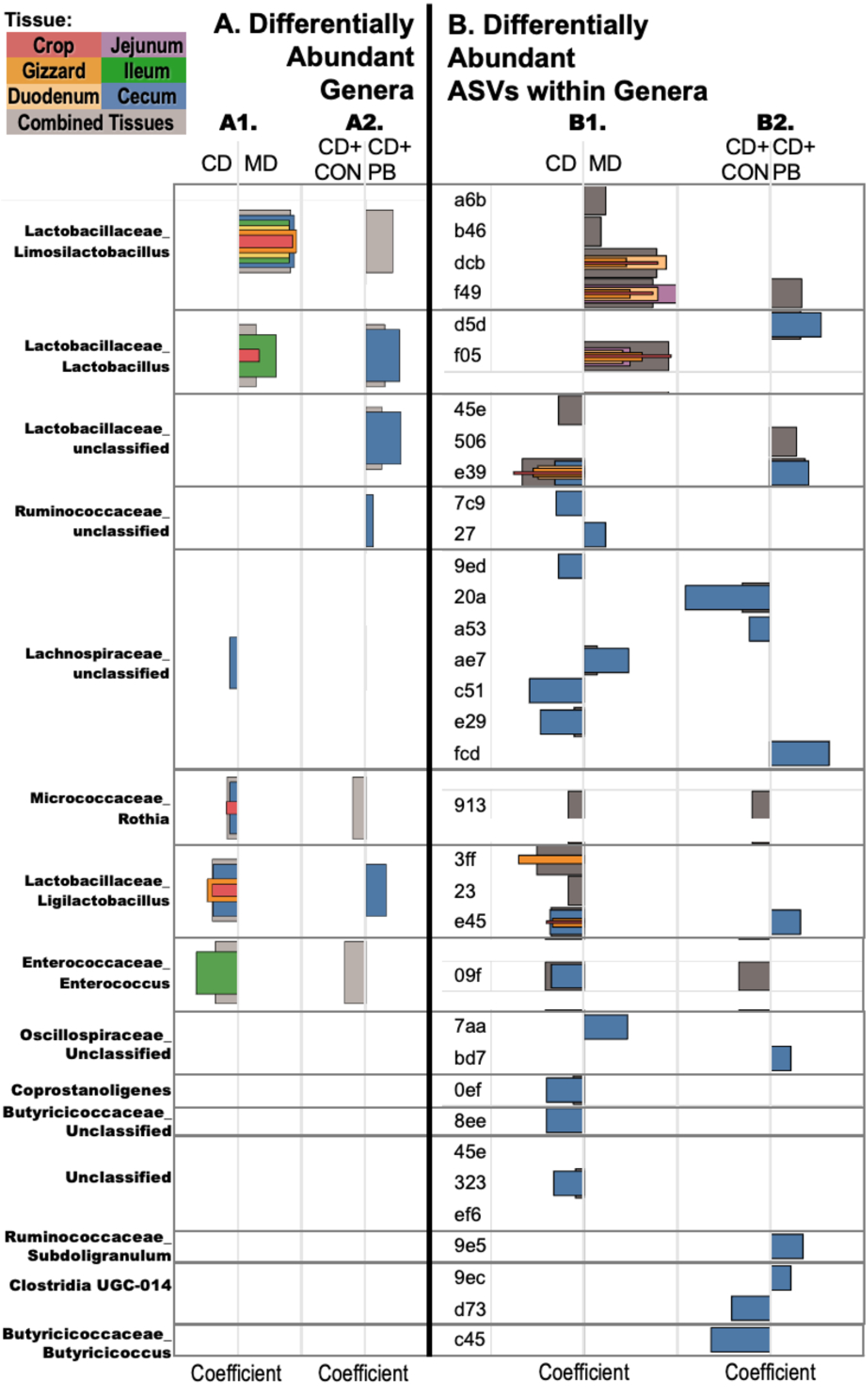
Lactobacillaceae taxa are central to differential abundance responses under both diet and PB treatment, with heterogeneity within genera at the ASV level. Differential abundance analysis was performed on bacterial genera (left panels) and constituent amplicon sequence variants (ASVs; right panels) in response to diet and probiotic treatment. Colored bars indicate the gastrointestinal region of origin: crop (red), gizzard (orange), duodenum (yellow), jejunum (purple), cecum (blue), and all tissues considered together (gray). Bars represent the average effect size (coefficient) from the differential abundance model for each genus or ASV. Only genera and ASVs that were significantly differentially abundant (FDR-adjusted p < 0.1) and related to PB supplementation are shown. Remaining data can be found at Supplemental Table 2. **(A1)** Genera that were significantly different in abundance between CD and MD, regardless of PB treatment. **(A2)** Genera that were significantly different in abundance between PB and CON birds within the CD. **(B1)** ASVs within the genera in (A1) that were significantly differentially abundant between basal diets. **(B2)** ASVs within the genera in (B1) that were significantly different between PB and CON treatments in CD-fed birds.

In contrast, there was stronger evidence for PB-induced microbial shifts in CD-fed birds, particularly in the cecum (Figure 2A2, Figure 2B2, Supplemental Figure 2). Genera enriched in CD+PB animals included *Lactobacillus*, *Rothia*, *Ligilactobacillus*, and an uncultured genus within the Lactobacillaceae family (Figure 4A2), each driven by ASVs within each genus (Figure 4B2). Although *Enterococcus* abundance appeared to be reduced with PB treatment (Figure 4A2, Figure 4B2), this effect was largely attributable to a single animal in the CD+CON group (data not shown). While high variability prevents a firm conclusion, the potential for PB to modulate *Enterococcus* warrants further investigation.

Basal diet also shaped background microbial communities in ways that likely influenced PB responsiveness. Several PB-associated taxa, including *Lactobacillus* and *Limosilactobacillus*, were more abundant in MD regardless of PB treatment (Figure 4A1, Figure 4B1), which may have reduced the detectability of PB-induced changes within MD. In contrast, *Ligilactobacillus* and its associated ASVs were more abundant in CD across multiple tissues (Figure 4A1, Figure 4A2), consistent with earlier clustering results (Figure 2A). Given that serum from CD-fed birds led to greater ATP production *in vitro* (Figure 3) compared to MD, *Ligilactobacillus* and its associated ASVs are stronger candidates for contributing to the immune energizing effect than taxa enriched in MD-fed animals.

Together, these results reveal diet-dependent shifts in microbial composition, with PB-associated changes far more apparent in CD than MD, despite the similar systemic energizing effect. These patterns suggest that more subtle microbial functional shifts, not evident in compositional profile, may vary within diet and treatment.

### Predicted microbial pathways related to mevalonate and carbohydrate metabolism are enriched in CD+PB ceca

To identify functional shifts, predicted metagenomic pathway abundances from 16S profiles were extracted via PICRUSt2. CD+PB birds exhibited enrichment in predicted microbial functions related to carbohydrate, mevalonate, and amino acid metabolism pathways, in the cecum (Figure 5a). In contrast, no significant differences were detected in the predicted functional profiles of MD-fed animals. The ASV contributing most strongly to the shifts in CD+PB was *Ligilactobacillus* e451, which had the highest proportional influence on 3 enriched pathways: the mevalonate pathway I, geranylgeranyl diphosphate (GGPP) biosynthesis via mevalonate, and the superpathway of uridine diphosphate glucose (UDP)-glucose-derived O-antigen building blocks (Figure 5B). Because *Ligilactobacillus* is gram-positive and does not synthesize O-antigen, this latter result is likely a classification artifact in the MetaCyc database. Instead, it is more likely that the predicted genes are involved in UDP-glucose enrichment, which shares many functional genes and aligns with *Ligilactobacillus* biology (28, 29).

**Figure 5.**
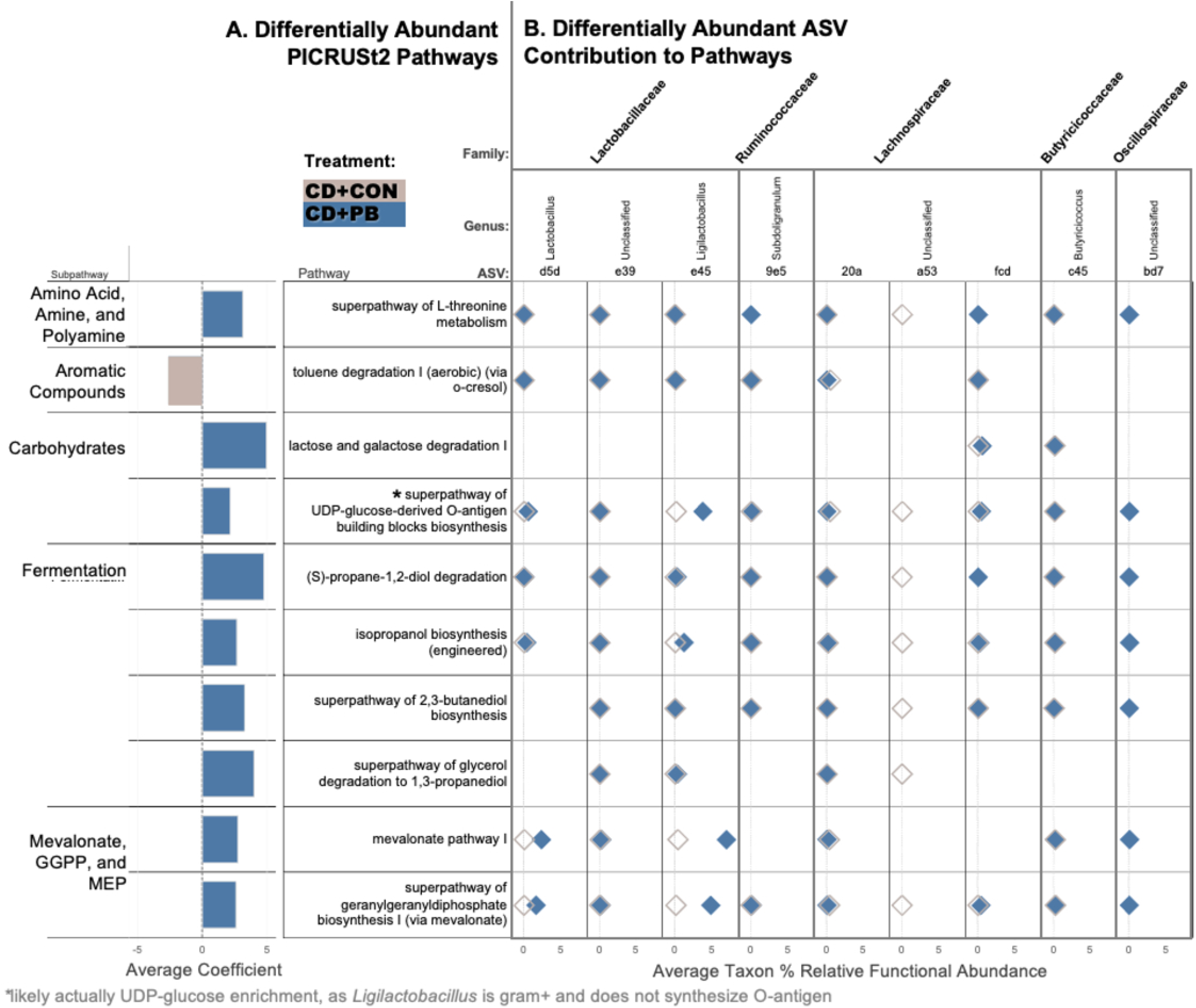
PB treatment enriches predicted fermentation, carbohydrate, and mevalonate-associated pathways in the cecum of CD birds. PICRUSt2 was used to infer microbial functional potential from 16S rRNA gene sequences in cecal samples from birds fed CD, comparing PB and CON groups. **(A)** Differentially abundant MetaCyc pathways (FDR < 0.1) between CD+PB and CD+CON cecal samples. Pathways are grouped by functional category (e.g., carbohydrate metabolism, fermentation, isoprenoid biosynthesis), and bars represent model-estimated effect sizes (average coefficients) from the differential abundance analysis. **(B)** Contributions of differentially abundant amplicon sequence variants (ASVs) to the predicted abundance of each pathway shown in (A). Each column represents an ASV, and each row corresponds to a pathway. Gray circles indicate the average relative functional contribution of that ASV in CON, and blue circles represent the contribution in the PB. The gray shading highlights the change in ASV’s functional contribution between treatments. Only ASVs contributing >0.1% to pathway abundance are shown. Circle size reflects the relative abundance of the ASV within cecal samples.

These predictions indicate an enrichment of genes associated with mevalonate pathway and isoprenoid biosynthesis in CD+PB birds. If reflected at the metabolite level, these changes could contribute to higher levels of serum isoprenoids or lipid intermediates (30, 31), molecules that immune cells can use as energetic substrates (32–34) or signaling mediators (35, 36). While speculative, these predicted functional differences suggest potential microbial pathway differences that warrant further investigation as contributors to the observed increase in immune cell ATP (Figure 3).

Microbial products such as isoprenoids, short-chain fatty acids, and amino acid derivatives have been shown in other species to modulate immune metabolism, membrane remodeling, and signaling cascades (37–40). While these pathways remain understudied in poultry, our findings suggest that diet-dependent microbial functions may contribute to systemic immunometabolic responses. Follow up experiments will test this hypothesis directly by quantifying serum and GI metabolites under probiotic treatment.

## Conclusion

Serum from PB-treated animals increased immune cell ATP production *in vitro* in both diet groups, despite minimal microbial community shifts in MD-fed animals. This unexpected disconnect indicates that detectable shifts in overall community structure are not strictly required for PB-associated physiological effects under the conditions tested. Dietary context further shaped baseline immunometabolic function in that CD serum-treated cells produced more ATP at baseline than MD-treated cells. However, cellular energizing response to PB-fed serum was similar in both diets, indicating that diet-driven energetic baseline differences did not interfere with PB-related metabolic or signaling inputs.

In CD-fed birds, the pronounced shifts in both microbial composition and predicted function, especially involving isoprenoid and sugar nucleotide biosynthesis, point to members of the Lactobacillaceae family, such as *Ligilactobacillus* and *Limosilactobacillus,* as prominent taxa in CD+PB animals that coincides with predicted functional shifts and enhanced immune cell ATP production. While these associations remain correlative, they support a testable hypothesis that PB influences serum lipid and isoprenoid composition, which in turn contributes to immune cell energetics.

More broadly, these results underscore that the absence of major structural change in the microbiota does not equate to the absence of physiological impact. Community profiling is a convenient, accessible, and widely used proxy for probiotic activity, but substantive host responses can occur without detectable shifts in community structure. Effective evaluation of probiotic interventions depends on integrating functional and host-response measures alongside compositional evaluation. Follow up work will quantify serum and GI metabolites under PB treatment to identify the microbial products driving the immune-energizing effect, supporting the broader goal of designing probiotic interventions guided by host outcomes rather than community composition alone.

## Data Availability

All raw 16S rRNA gene sequencing reads generated in this study have been deposited in the NCBI Sequence Read Archive (SRA) under BioProject accession PRJNA1450620 and are publicly available at https://www.ncbi.nlm.nih.gov/bioproject/PRJNA1450620. Sample metadata associated with each sequence file are included in the BioProject submission. Analytical scripts used in this study are available from the corresponding author upon reasonable request. All other data supporting the conclusions of this article are included within the article and its supplementary files.

## Ethics Approval

All animal procedures were approved by the North Carolina State University Institutional Animal Care and Use Committee (OLAW #A3331-01)

